# A single dsRNA spray silences *VAMT* and shifts habanero pepper fruit metabolism towards capsinoids

**DOI:** 10.64898/2026.09.01.748336

**Authors:** Estefania Arellano-Ordoñez, Amanda Kim Rico-Chávez, Irineo Torres-Pachecho, Rosalia Virginia Ocampo-Velázquez, Ramón Gerardo Guevara-González, Christopher Cedillo

## Abstract

Capsaicinoids are synthesized in the placenta of *Capsicum* fruit, where vanillylamine aminotransferase (VAMT) catalyzes the formation of vanillylamine, the precursor of the pathway. The modulation of pungency has relied on genetic breeding and transgenic approaches, and this pathway has not been addressed by spray-induced gene silencing. The aim of this study was to evaluate whether a single non-invasive spray of double-stranded RNA (dsRNA) targeting *VAMT* allows the gene to be silenced and capsaicinoid accumulation to be modified in *Capsicum chinense* fruit. The molecule was designed *in silico* and applied at 10 days post-anthesis. Pedicel injection reduced the *VAMT* transcript in a dose-dependent manner, with three levels of inhibition distinguishable from one another. Spraying with surfactant reduced it by 86.2 %, a magnitude statistically indistinguishable from the 90.6 % obtained by injection, and also reduced the *Pun1* transcript, a co-regulation previously described only as a difference between cultivars. Analysis by gas chromatography coupled to mass spectrometry showed reductions of 84.4 % in capsaicin and 68.5 % in dihydrocapsaicin, the loss of nonivamide and one further vanillylamine-derived compound, and the detection of capsiate and a second capsinoid, absent in control fruits. The siRNA was detected in non-treated tissues, and a single topical application is therefore sufficient to silence an endogenous biosynthetic gene and shift the metabolic profile of the fruit without genetic modification.

**Key Message:** *VAMT* was silenced by a single spray as effectively as by injection, capsaicin was reduced by 84.4 %, *Pun1* was co-reduced, and the siRNA was detected in non-treated tissues

## Introduction

Spray-induced gene silencing (SIGS) is a non-transgenic approach that relies on the topical application of double-stranded RNA (dsRNA) to trigger the post-transcriptional silencing of specific genes (Koch et al., 2016). The RNAi mechanism involves the processing of dsRNA by Dicer into small interfering RNAs (siRNAs), which are loaded into the RNA-induced silencing complex to degrade complementary mRNA transcripts. It has been shown that naked dsRNA can be internalized and processed by the RNA interference (RNAi) machinery of plants without the need for carriers (Nityagovsky et al., 2022), and that siRNAs can move systemically through the phloem and silence genes in distal tissues (Cisneros et al., 2022).

These properties have made SIGS a versatile tool, employed from pathogen control to the modulation of specialized metabolism. Foliar application of dsRNA against MYB repressors in tomato increased anthocyanin accumulation (Suprun et al., 2023) and external application of dsRNA reduced the expression of anthocyanin biosynthesis genes in *Arabidopsis thaliana* (Kiselev et al., 2021a). In edible tissue, postharvest spraying of dsRNA reduced the enzymatic browning of fresh-cut potato (Chen et al., 2023), whereas silencing of sugar transporters modified the interaction of the plant with nematodes (Warnock et al., 2016).

In chili pepper (*Capsicum* spp.), the compound of greatest interest is capsaicin, which is responsible for pungency and has several pharmaceutical applications (Reddy et al., 2024). Its biosynthesis converges on the enzyme *VAMT* (vanillylamine aminotransferase), which forms vanillylamine from vanillin; its biochemical characterization led to the previous designation of putative aminotransferase (*pAMT*) being replaced by *VAMT* (Nakaniwa et al., 2024). *VAMT* occupies a branch point of the pathway: in its absence, the flux is diverted towards the production of capsinoids, which are pungency-free analogues of capsaicinoids (Lang et al., 2009; Tanaka, 2025). Its expression is predominantly placental and correlates with capsaicin accumulation during fruit development (Arce-Rodríguez & Ochoa-Alejo, 2019). These features make *VAMT* a suitable target for modulating pungency by a non-transgenic route.

This pathway, however, has not been addressed by SIGS. The aim of this study was therefore to evaluate whether a single non-invasive spray of dsRNA targeting VAMT silences the gene and modifies capsaicinoid accumulation in *Capsicum chinense* fruit, comparing spraying with pedicel injection and examining whether the siRNA reaches non-treated tissues. The results show that topical delivery matches injection in the inhibition of the gene, that the blockade of the node shifts the metabolic profile of the fruit towards capsinoids, and that the siRNA moves beyond the application site.

## Materials and Methods

### Plant material and growth conditions

Certified habanero pepper seeds (*Capsicum chinense* Jacq.) cv. “El Jaguar” were germinated in 12-cavity trays (10 × 12 × 18 cm) filled with a 3:1 (v/v) mixture of commercial peat and perlite. Upon reaching the four-true-leaf stage, seedlings were transplanted into 20 L greenhouse bags containing the same substrate mixture. The experiments were conducted in experimental greenhouse no. 6 (36 m²) at the Amazcala Campus of the Faculty of Engineering, Universidad Autónoma de Querétaro. The greenhouse was naturally ventilated through manually operated windows and equipped with an automated drip irrigation system with pressure-compensating emitters. The injection experiment was carried out in November 2024 and the spray experiment in September 2025. Nutrients were supplied by fertigation according to the formulation reported in Table S1 of the Supplementary Information. Irrigation was delivered in eight daily events for a total of 400 mL per plant per day.

### Developmental time-course sampling

For the analysis of *VAMT* transcript expression during fruit development, twelve habanero pepper plants were used. Flower buds at anthesis were individually tagged so that the fruit stage could be recorded. Fruits derived from tagged flowers were harvested at 0, 3, 5, 10, 20, 30, and 40 DPA, so that three biological replicates of one-fruit experimental units were obtained per stage. Whole fruits were processed for the 0 to 10 DPA stages, whereas for the 20 to 40 DPA stages, only placental tissue was dissected and processed. In addition, one leaf and one unpollinated flower were collected from each of three randomly selected plants as reference tissues, in order to assess the presence of the *VAMT* transcript in organs other than the developing fruit.

### In silico design and synthesis of dsR*VAMT*

The small RNA molecule targeting *VAMT* was designed according to Cedillo-Jiménez et al. (2024), using as target the *VAMT* coding sequence deposited under NCBI Reference Sequence NM_001324706.1. The design was based on endogenous microRNAs with potential complementarity to the *VAMT* transcript. These were predicted with psRNAtarget V2 (Dai et al., 2018), interrogating the published *Solanum lycopersicum* miRNA dataset, selected for its phylogenetic proximity to *Capsicum* within the Solanaceae. The closest candidate miRNA sequence was optimized so that full complementarity with the target region was achieved, giving rise to siR*VAMT*. Target site accessibility (ΔG open) and hybridization energy (ΔG hybrid) between siR*VAMT* and the *VAMT* transcript were subsequently calculated with RNAup 2.5.1 (ViennaRNA Web Services) using default parameters (Mückstein et al., 2006).

The siR*VAMT* strand (UUCACAAACUCUGUAGAAAGU) and its complementary strand (AUUCUACAGAGUUUGUGAAUU) were each synthesized carrying a 5′ phosphorothioate linkage and a 3′ 2′-O-methyl ribonucleotide. These modifications were incorporated so that molecular stability and nuclease resistance would be increased, as previously described (Szabat & Kierzek, 2017). The complementary strand was designed in such a way that two-nucleotide 3′ overhangs were left on both strands, so that duplex formation would be facilitated, as previously described (Ghosh et al., 2009). The oligonucleotides were synthesized by T4 Oligo (Irapuato, Mexico).

In order to evaluate potential off-target interactions, a target prediction analysis was performed with psRNAtarget V2 (Dai et al., 2018) against the *Capsicum annuum* transcriptome (CDS, v2), given that no *Capsicum chinense* library is available on the platform. Predictions were interpreted under an Expectation threshold of ≤ 2.0, based on the balance between sensitivity and specificity reported for the tool (Lück et al., 2019).

### Preparation and application of dsR*VAMT*

dsR*VAMT* was generated by annealing equimolar amounts (1:1) of the siR*VAMT* strand and its complementary strand, followed by incubation at 90 °C for 1 min and gradual cooling to room temperature, according to Dubrovina et al. (2020). The applied molecule was therefore a short dsRNA of 21 nt per strand. siRVAMT designates the guide strand expected to direct silencing, whereas dsRVAMT designates the double-stranded molecule that was actually applied.

For the injection treatments, 5 µL of dsR*VAMT* solution were administered into the fruit pedicel with a sterile insulin syringe, one fruit per plant, at 10 DPA. Three doses were evaluated: 1000 pmol, 100 pmol, and 10 pmol. Control fruits were injected with an equal volume of sterile RNase-free distilled water. Each treatment comprised three biological replicates, with one plant defined as the experimental unit, and plants were arranged in a completely randomized design.

For the spray treatment, 1000 pmol of dsR*VAMT* contained in 5 µL, matching the most concentrated injection treatment, were mixed with 100 µL of the surfactant EcoNano MegaKlinner (AGRONATURALIA; manufacturer’s technical data sheet in Table S2) and brought to 5 mL with sterile nuclease-free water, giving a final surfactant concentration of 2 % (v/v). The controls comprised fruits sprayed with water, fruits sprayed with 2 % (v/v) surfactant in distilled water, and a positive inhibition control consisting of injected dsR*VAMT* (1000 pmol). Treatments were applied at 10 DPA, at 08:00 h. Each treatment comprised three biological replicates (one plant as the experimental unit) in a completely randomized design, with treatments spatially separated so that aerosols from different treatments could not come into contact.

### RNA extraction

Approximately 100 mg of placental tissue were collected at 20 DPA (10 days post-treatment), frozen in liquid nitrogen, and ground to a fine powder with mortar and pestle. Total RNA was extracted with TRIzol™ reagent (Ambion, Thermo Fisher Scientific) following the manufacturer’s instructions, and the RNA pellet was resuspended in 30 µL of sterile RNase-free distilled water. Total RNA concentration was subsequently adjusted to 500 ng/µL for the developmental time-course analysis and to 230 ng/µL for the dsRNA treatment experiments, in which lower RNA yields were obtained.

### Semi-quantitative endpoint RT-PCR

First-strand cDNA synthesis was performed with the RevertAid kit (Thermo Fisher Scientific) following the manufacturer’s instructions, using 230 ng of total RNA per reaction for the treatment experiments and 500 ng for the developmental time course. Reverse transcription was carried out in a Techne TC-3000G thermocycler: samples were incubated at 65 °C for 5 min and held on ice for 2 min, followed by 5 min at 25 °C, 60 min at 42°C, 5 min at 70 °C, and a final hold at 4 °C.

Transcript abundance was assessed by semi-quantitative endpoint RT-PCR followed by densitometric analysis. Amplification was performed with the Maxima SYBR Green kit (Thermo Fisher Scientific) under the conditions specified by the supplier. Amplification was carried out in the same thermocycler with an initial denaturation at 95 °C for 10 min, followed by 38 cycles of 95 °C for 15 s, annealing for 30 s, and 72 °C for 30 s, with a final hold at 4 °C. Gene-specific annealing temperatures were 60 °C for VAMT, 61.4 °C for Pun1, and 62.9 °C for β-actin.

Expected amplicon sizes were verified prior to analysis: 210 bp for *VAMT* (LC423555), 126 bp for *Pun1* (LC423556), and 198 bp for *β-actin* (reference gene; AY572427). The primers, taken from Sano et al. (2022), are reported in Table S3. β-actin was selected as the reference gene because it has been employed as an internal control in expression analyses of capsaicinoid biosynthetic genes in placental tissue of Capsicum throughout fruit development (Zhang et al., 2016).

Gel images were acquired with a Gel Doc™ EZ System (Bio-Rad) and exported with ImageLab™ software (Bio-Rad, version 6.0.1). Densitometric analysis was subsequently performed in ImageJ v1.54r: images were converted to 8-bit grayscale, background signal was measured adjacent to each band, and background subtraction was performed with the Subtract function. Relative transcript levels were calculated by normalizing the area under the curve of *VAMT*, or *Pun1* against that of the *β-actin* reference gene for each sample.

### Systemic siR*VAMT* detection by stem-loop RT-PCR

The presence or absence of translocated siRVAMT was detected by specific reverse transcription with stem-loop oligonucleotides followed by endpoint PCR, based on the stem-loop method of Varkonyi-Gasic et al. (2007): reverse transcription was performed with a stem-loop primer (5′-GTTGGCTCGGTGCAGGGTCCGAGGTATTCGCACCAGAGCCAC-3′; Table S3) in a Techne TC-3000G thermocycler with an initial incubation at 16 °C for 30 min, followed by 60 cycles of 30 °C for 30 s, 42 ° C for 30 s, and 50 °C for 1 s, and a final incubation at 85 °C for 5 min in order to inactivate the reverse transcriptase.

The cDNA was subsequently amplified by endpoint PCR with a specific forward primer (5′-TTCACAAACTCTGTAGAAAG-3′) and a universal reverse primer (5′-GTGCAGGGTCCGAGGT-3′; Table S3) in a Techne TC-3000G thermocycler, with initial denaturation at 95 °C for 5 min, followed by 40 cycles of 95 °C for 15 s and 60 °C for 1 min. PCR products were visualized with a Gel Doc™ EZ System (Bio-Rad) and analyzed with ImageLab™ software (Bio-Rad, version 6.0.1) after electrophoresis under the conditions described above. Detection was recorded as presence or absence of band.

### Quantification of capsaicinoids and capsinoids by GC-MS

GC-MS analysis was performed on control fruits and on fruits treated with dsRVAMT by spraying, with three biological replicates per group. Placental tissue was collected from each fruit at 20 DPA (10 days post-treatment), lyophilized, and ground. Subsequently, 500 mg of the resulting powder were extracted with acetonitrile (1:20, w/v) by sonication at 30 °C for 3 h, followed by orbital shaking at 200 rpm for 12 h. Samples were centrifuged at 12,000 ×g and 10 °C for 10 min. Extraction was repeated with acetonitrile (1:10, w/v) and the combined supernatants were evaporated under a stream of nitrogen.

Dry extracts were resuspended in 200 µL of acetonitrile, centrifuged again, and analysed by gas chromatography coupled to mass spectrometry (GC-MS) on an Agilent 5975C system operated in simultaneous scan/SIM mode with electron ionization at 70 eV. Separation was performed on an HP-5MS column (30 m × 250 µm × 0.25 µm) with helium as carrier gas at a constant flow of 1.2 mL/min, and 1 µL was injected in splitless mode with the inlet at 250 °C. The oven program was: 90 °C held for 1 min, ramped at 12 °C/min to 210 °C (held 1 min), then at 4 °C/min to 290 °C (held 1 min), for a total run time of 33 min. The transfer line, ion source, and quadrupole were maintained at 250, 230, and 150 °C, respectively, with a solvent delay of 3 min and a scan range of m/z 29–500.

Compound identities were assigned by comparison of full-scan spectra with the NIST 11 library and by retention-time matching against capsaicin, dihydrocapsaicin, and capsiate standards. Quantification was performed in SIM mode, with m/z 137, the vanillyl fragment shared by capsaicinoids and their esters, used as the quantifier ion, and co-eluting analytes were resolved by their retention times. Capsaicin (Sigma-Aldrich, ≥ 95 %, cat. no. M2028), dihydrocapsaicin (Sigma-Aldrich, ≥ 85 %, cat. no. M1022), and capsiate (Sigma-Aldrich, analytical standard, ≥ 90 %, cat. no. 90526) were used to construct their respective calibration curves. Compounds for which no authentic standard was available were semi-quantified against the curve of the structurally closest standard, in accordance with the criteria for surrogate-standard semi-quantification set out by Malm et al. (2021).

Calibration ranges, coefficients of determination, and limits of detection and quantification of each analyte are reported in Table S4. Capsaicinoids and capsinoids were extracted from the placenta of each fruit, and each extract was injected once; three biological replicates were analyzed per treatment, and technical replicates were not performed.

### Statistical analysis

Statistical analyses were performed with GraphPad Prism version 8 (GraphPad Software, Boston, MA, USA). Transcript abundance data were evaluated by one-way analysis of variance (ANOVA) at α = 0.05, followed by Tukey’s post hoc multiple comparison test when significant differences were detected (p < 0.05). Metabolite concentrations, for which two groups were compared, were evaluated by an unpaired two-tailed Student’s t-test at α = 0.05. Normality of residuals (Shapiro-Wilk and Anderson-Darling) and homogeneity of variances (Brown-Forsythe) were verified prior to each analysis. Compounds recorded as not detected were assigned a value of zero for the calculation; those for which an entire group was not detected were not subjected to inferential testing and are reported as detection frequency. The summed concentration of the five vanillylamine-derived amides was obtained by adding the individual concentrations of each replicate and averaging per group; compounds recorded as not detected were included with a value of zero. Three biological replicates were analysed per treatment, and one plant was defined as the experimental unit.

## RESULTS

### 1. Abundance of *VAMT* transcripts during fruit development

In order to determine the optimal stage for dsRNA application, *VAMT* transcript abundance was monitored throughout fruit development at 0, 3, 5, 10, 20, 30, and 40 days post-anthesis (DPA; Fig. 1A). Since the placenta cannot be reliably dissected from fruits smaller than 2 cm, the 0 to 10 DPA stages were processed as whole fruit and the 20 to 40 DPA stages as placental tissue, so that the two intervals are presented separately.

**Fig. 1.**
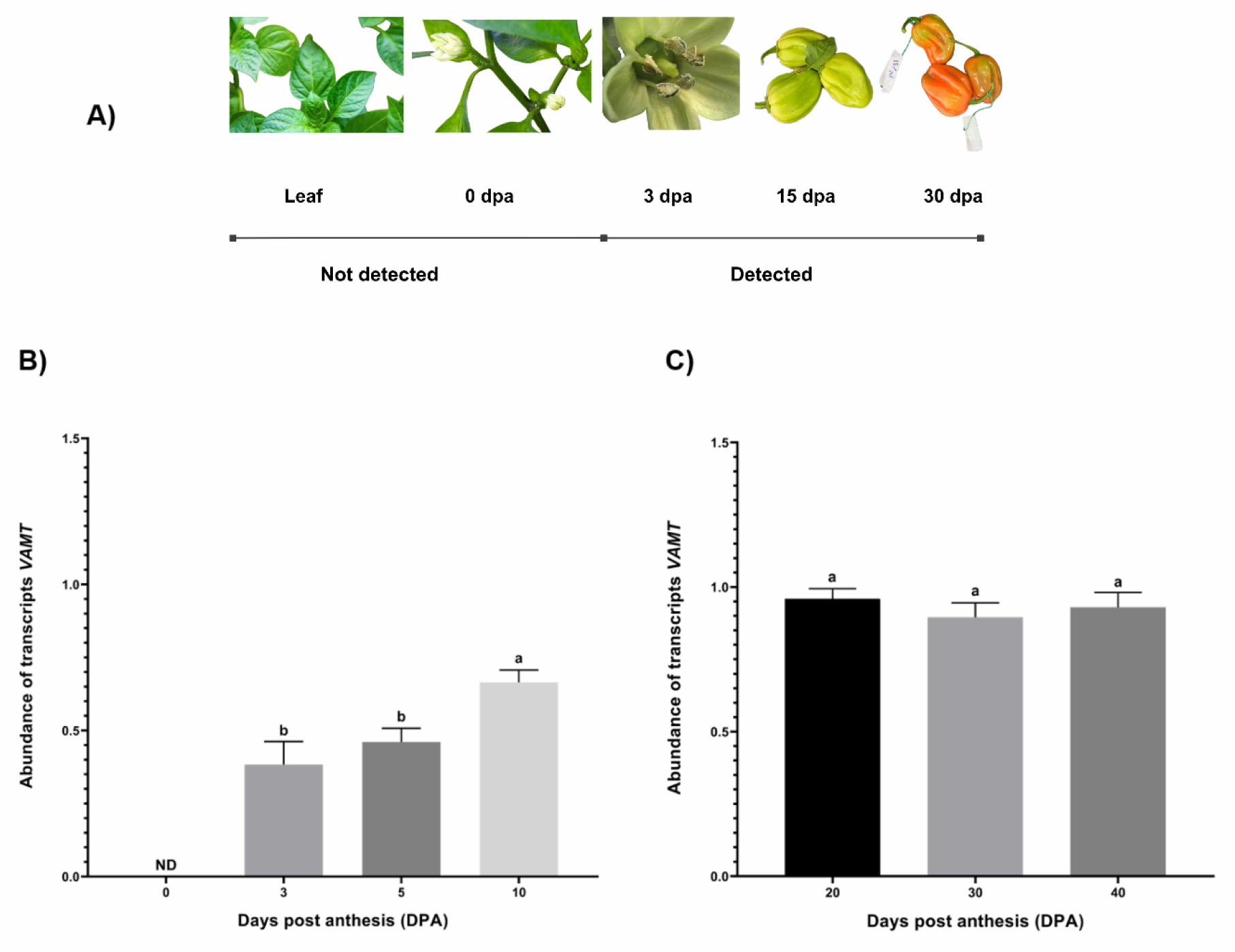
*VAMT* expression in different tissues and developmental stages. **A** Representative stages of plant and fruit development. **B** Abundance of *VAMT* transcript at 0, 3, 5, and 10 DPA, determined in whole fruit. **C** Abundance of *VAMT* transcript at 20, 30, and 40 DPA, determined in placental tissue. Values represent mean ± SD of three biological replicates. Different letters indicate significant differences among stages within each panel (p < 0.05, Tukey’s test). Samples were normalized to *β-actin*. ND, *VAMT* not detected Based on the temporal expression pattern, the 10 DPA stage was selected as the point of dsRNA application, since it occurs after transcript detection and before *VAMT* transcript levels stabilize at higher values during the later developmental stages.

The *VAMT* transcript was not detected in leaves or in unpollinated flowers. In whole fruit, it was not detected at 0 DPA but became detectable from 3 DPA onwards, and its abundance showed significant differences among the stages in which the transcript was detected (one-way ANOVA, F(2,6) = 18.60; p = 0.0027), with equivalent levels at 3 and 5 DPA and an increase at 10 DPA (Fig. 1B). In placental tissue, abundance did not differ among 20, 30, and 40 DPA (F(2,6) = 1.44; p = 0.308), which suggests that expression stabilizes from 20 DPA onwards (Fig. 1C).

### 2. Design of interfering dsRNA targeting *VAMT*

A computational strategy was employed in order to design an RNAi molecule directed against *VAMT*. Target regions were identified by in silico prediction of miRNA and target interaction against the *VAMT* transcript. Among the candidates, sly-miR9472-3p from tomato was selected on the basis of its complementarity to a region spanning nucleotides 1131 to 1151 of the *VAMT* transcript and on the basis of its phylogenetic proximity, since pepper and tomato both belong to the Solanaceae. Since mismatches were present in the interaction, the sequence was optimized until full complementarity was achieved, and the resulting molecule was designated siRVAMT (UUCACAAACUCUGUAGAAAGU; Fig. 2A).

**Fig. 2.**
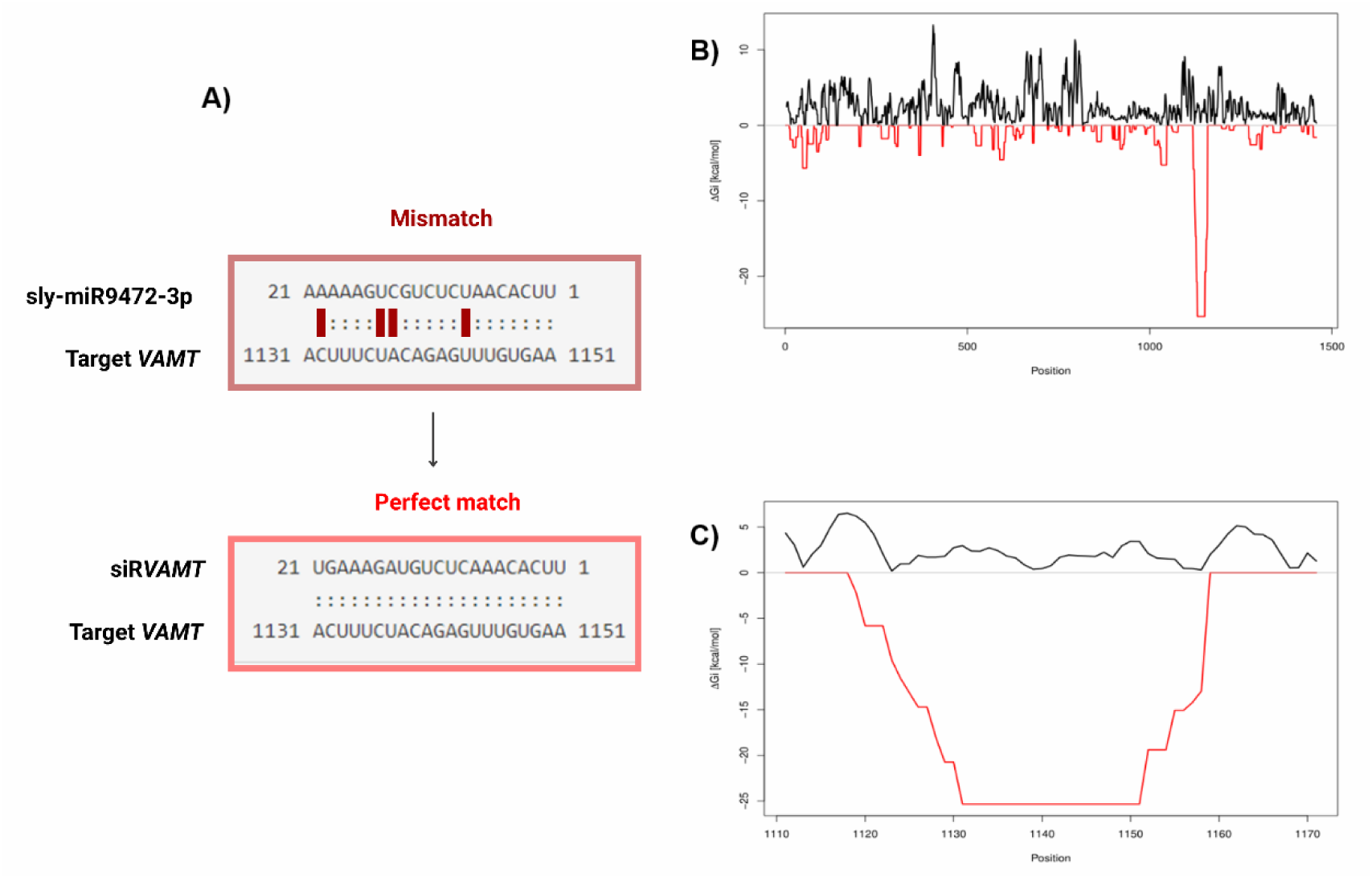
Design basis for an RNAi molecule targeting *VAMT*. **A** Alignment of the endogenous sly-miR9472-3p with the *VAMT* transcript showing partial complementarity (mismatch), and the optimized siR*VAMT* sequence displaying full complementarity (perfect match) to the target region (nt 1131 to 1151). **B** Genome-wide thermodynamic profile of *VAMT*. The red line (siR*VAMT*) highlights a highly favorable hybridization energy peak (ΔG approximately -25.32 kcal/mol) at the selected target site, whereas the endogenous miRNA (black line) shows weaker interaction across the transcript. **C** Local target site accessibility analysis. The red profile represents the optimized siR*VAMT*, showing a marked decrease in ΔG (opening energy), indicating higher accessibility of the target region compared to the endogenous miRNA (black line)

Thermodynamic analysis indicated that the selected target region exhibited low structural accessibility energy (ΔG open = 1.42 kcal/mol), suggesting that little energy is required for the binding site to be exposed. In addition, the hybridization energy between *VAMT* and siR*VAMT* was found to be highly favorable (ΔG hybrid = −25.32 kcal/mol), which supports a stable interaction (Fig. 2B–C).

The specificity of siR*VAMT* was evaluated by predicting potential off-targets. Under the adopted threshold, which has been reported to recover 82.6 % of experimentally validated targets at a false-positive rate of 26.9 % (Dai and Zhao, 2011), no transcript other than VAMT was retrieved. Once specificity had been established *in silico*, the silencing capacity of dsR*VAMT* was evaluated through the route that offers the greatest assurance of internalization, that is, by injection.

### 3. dsRVAMT injection reduces the abundance of VAMT and Pun1 transcripts in a dose-dependent manner

In order to determine whether VAMT silencing could be achieved once uptake barriers had been bypassed, dsRVAMT was delivered by direct injection at three doses. Significant differences among treatments were detected for the VAMT target transcript (one-way ANOVA, F(3,8) = 642.82; p < 0.0001). A mean abundance of 0.7380 ± 0.0171 was exhibited by the control group, and transcript levels were significantly reduced by all three dsRNA treatments. The most pronounced reduction was observed at 1000 pmol dsRVAMT, with a mean of 0.0775 ± 0.0120, which represents an 89.5 % decrease (p < 0.0001). Significant reductions of 36.2 % and 26.0 % were also recorded at 100 pmol (0.4706 ± 0.0285) and 10 pmol (0.5462 ± 0.0138), respectively (p < 0.0001 in both cases; Fig. 3).

**Fig. 3.**
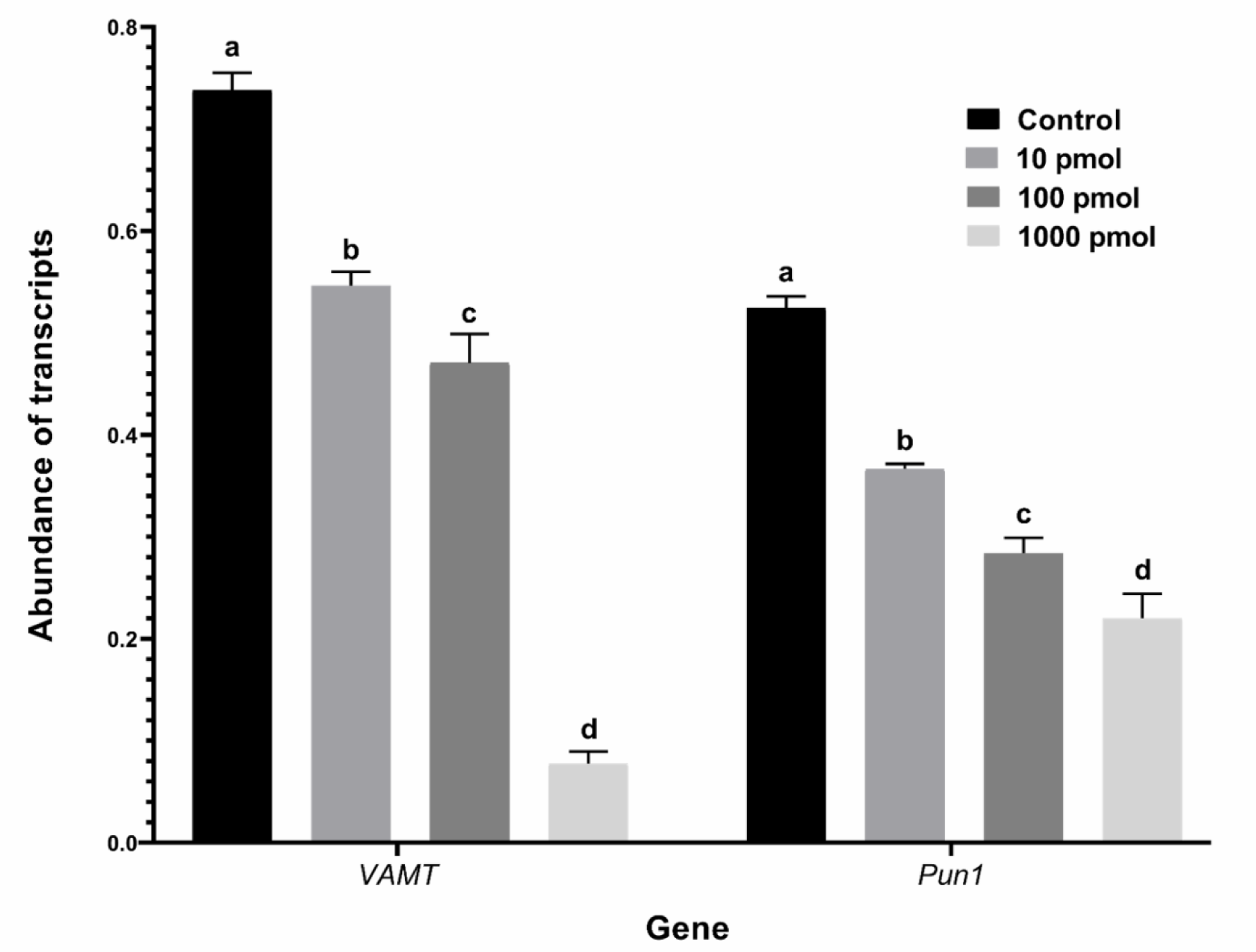
Abundance of *VAMT* and *Pun1* transcripts after dsRNA injection. Values represent mean ± SD of three biological replicates. Different letters indicate significant differences among treatments within each gene (p < 0.05, Tukey’s test). Samples were normalized to *β-actin*

When compared, the three doses differed significantly from one another. Taken together, these results indicate that VAMT transcript abundance followed a dose-dependent relationship across the three levels evaluated, and that silencing is achieved when uptake barriers are bypassed.

The abundance of the *Pun1* transcript, which encodes the acyltransferase that condenses vanillylamine with the fatty acid chain, was evaluated in the same samples. Significant differences among treatments were likewise detected (one-way ANOVA, F(3,8) = 222.01; p < 0.0001). A mean abundance of 0.5246 ± 0.0112 was exhibited by the control group, and transcript levels were significantly reduced by the three doses: 58.0 % at 1000 pmol (0.2203 ± 0.0238), 45.9 % at 100 pmol (0.2841 ± 0.0149), and 30.1 % at 10 pmol (0.3668 ± 0.0048), with p < 0.0001 in all three cases (Fig. 3). As for *VAMT*, the three doses were found to differ significantly from one another. *Pun1* abundance therefore decreased in a dose-dependent manner and in the same order as that of *VAMT*, although the magnitude of the reduction was smaller at each level.

Since silencing had been demonstrated at the site of application, whether the silencing signal was able to move beyond that site was examined next.

### 4. Systemic detection of siR*VAMT* in distal non-treated tissues

In order to assess whether siR*VAMT* could be detected beyond the application site, distal non-treated tissues were analysed from plants that had received the 1000 pmol dsRNA injection. *siRVAMT* was detected in non-treated flowers and leaves from treated plants, whereas it was not detected in the same tissues from untreated plants. A dsRVAMT-injected fruit, obtained from an independent plant, was included as a positive amplification control (Fig. 4).

**Fig. 4.**
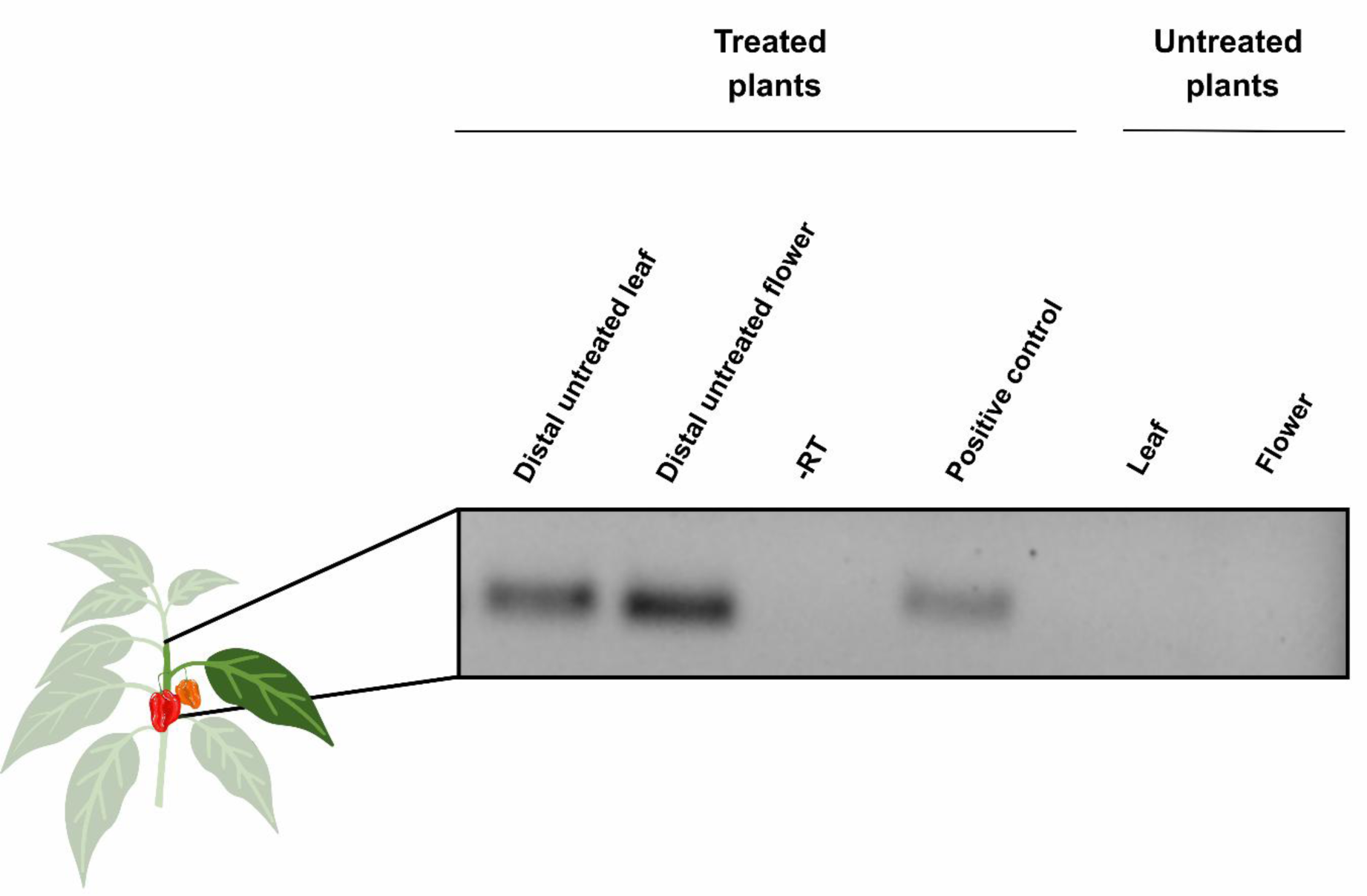
Detection of siR*VAMT* in distal tissues of treated plants. A 55 bp product corresponding to siR*VAMT*-derived siRNA was detected in non-treated leaves and flowers of dsRVAMT-treated plants. No amplification was observed in any tissue from untreated plants. A dsRVAMT-injected fruit from an independent plant was used as a positive control. The –RT control, corresponding to a treated fruit processed without reverse transcriptase, showed no amplification. Each lane corresponds to a representative sample from three biological replicates

### 5. Transcriptomic and metabolic responses associated with SIGS-mediated dsR*VAMT* application

The inhibitory effect of dsR*VAMT* delivered via SIGS was evaluated using a surfactant as the suspension medium. Statistical analysis showed significant differences in *VAMT* transcript abundance among treatment groups (one-way ANOVA, F(3,8) = 245.22; p < 0.0001). The water-sprayed control group had a mean of 0.7112 ± 0.0626, while the surfactant-only treatment (0.6971 ± 0.0426) did not differ significantly from that control (p = 0.9702). In contrast, both dsRNA applied by spraying with surfactant (0.0981 ± 0.0086) and injected dsRNA (0.0669 ± 0.0225) significantly reduced transcript abundance compared to the control (p < 0.0001 in both). These decreases corresponded to reductions of 86.2 % and 90.6 %, respectively. No significant differences were observed between the two dsRNA delivery methods (p = 0.7741; Fig. 5A).

**Fig. 5.**
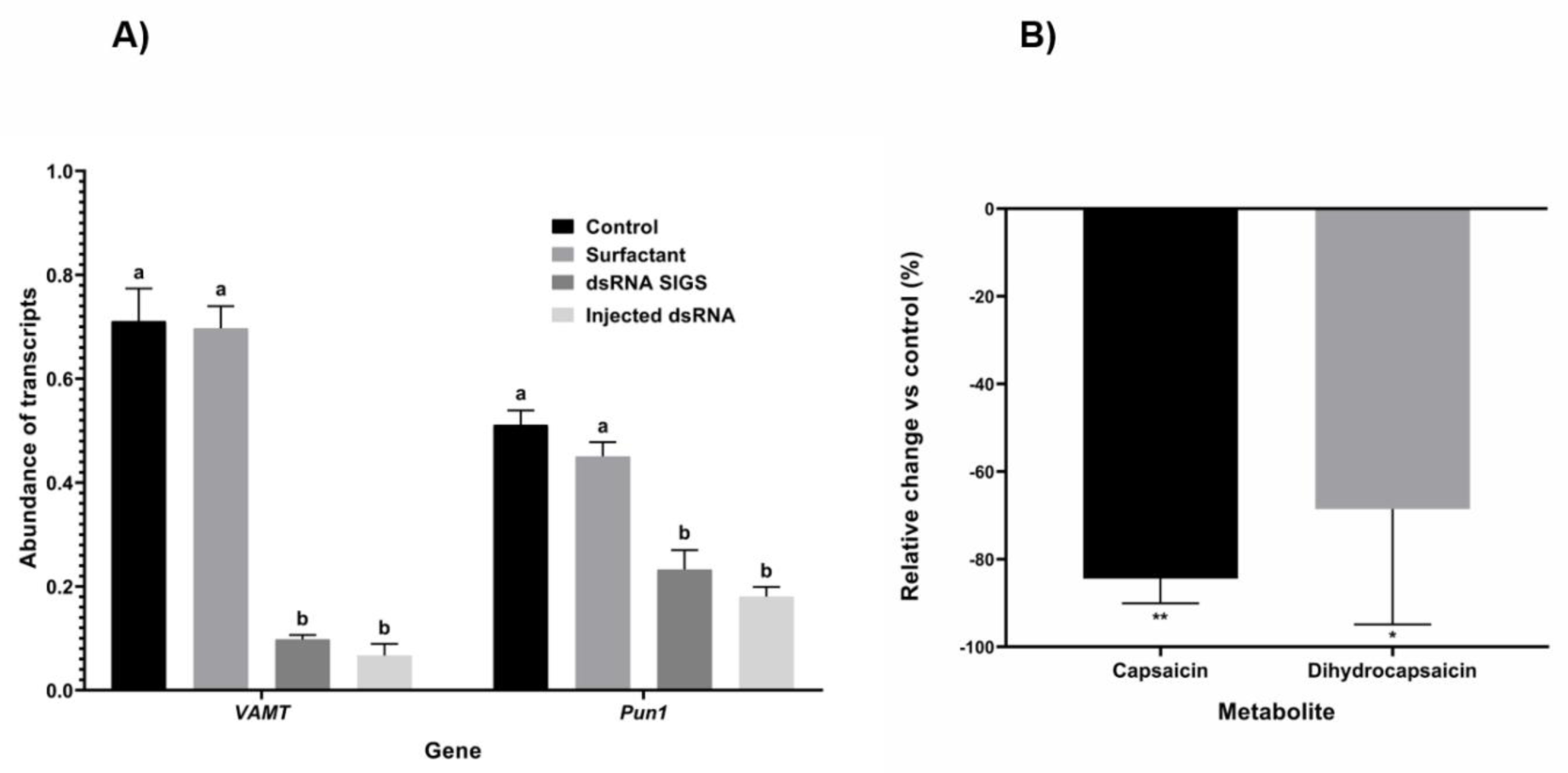
Transcriptomic and metabolic responses associated with SIGS-mediated dsRVAMT application. **A** Abundance of *VAMT* and *Pun1* transcripts in water-sprayed control fruits and after treatment with surfactant, dsRVAMT SIGS, and injected dsRVAMT. Different letters indicate significant differences among treatments within each gene (p < 0.05, Tukey’s test); samples were normalized to β-actin. **B** Relative change in the concentration of capsaicin and dihydrocapsaicin in dsRVAMT SIGS-treated fruits with respect to the control, expressed as a percentage. Values represent mean ± SD of three biological replicates. * p < 0.05, ** p < 0.01; unpaired two-tailed Student’s t-test

Regarding *Pun1*, the control group had a transcript abundance of 0.5111 ± 0.0277. The surfactant-only treatment (0.4508 ± 0.0270) did not differ significantly from the control (p = 0.1168). Both dsRNA treatments significantly reduced the abundance of *Pun1* transcripts: spray application with surfactant reduced it to 0.2329 ± 0.0372 (54.4 % reduction; p < 0.0001), and injected dsRNA reduced it to 0.1804 ± 0.0187 (64.7 %; p < 0.0001). No significant differences were observed between the two delivery methods (p = 0.1862; Fig. 5A).

In order to determine whether *VAMT* inhibition was associated with metabolic changes, the capsaicinoid and capsinoid profile of placental tissue was analysed by GC-MS in control fruits and in fruits sprayed with dsR*VAMT*.

Among the vanillylamine-derived amides, capsaicin was reduced by 84.4 % relative to the control (t = 6.221; p = 0.0034) and dihydrocapsaicin by 68.5 % (t = 3.463; p = 0.0257; Fig. 5B). Homodihydrocapsaicin, semi-quantified against the dihydrocapsaicin curve, was decreased by 58.2 %, a reduction that did not reach statistical significance (t = 1.529; p = 0.2011). Nonivamide was detected in the three replicates of the control group and in none of the treated replicates. An additional putative amide, eluting at 21.1 min and not identified, shared the m/z 137 quantifier ion and followed the same pattern: it was detected in the three control replicates and in none of the treated ones. The summed mean concentration of the five vanillylamine-derived amides was reduced by 81.9 % relative to the control (Table S5).

Conversely, two capsinoid esters absent from the control group were detected in the treated group. Capsiate was not detected in any of the three control replicates and was detected in two of the three treated replicates. An additional capsinoid, eluting at 17.5 min, followed the same pattern: it was not detected in the three control replicates and was detected in two of the three treated ones. Since the calibration curve obtained for capsiate reached a coefficient of determination of r² = 0.8994, below the r² ≥ 0.99 adopted as the acceptance criterion for quantification and met by the capsaicin and dihydrocapsaicin curves, both esters are reported as detection frequency and not as concentration. The compounds for which an entire group was non-detectable were not subjected to statistical testing. The limits of detection and quantification of all analytes are reported in Table S4, and the individual values in Table S5.

## Discussion

### 1. *VAMT* detection from 3 DPA

*VAMT* expression reaches its maximum at intermediate stages of fruit development, between 20 and 40 days after pollination, and declines during ripening (Aza-González et al., 2011). In the present work, the transcript was detected from 3 DPA, with increasing levels towards 10 DPA and stable levels from 20 DPA onwards. Most expression studies begin their sampling at 10 DPA (Ogawa et al., 2015), so that the interval between anthesis and that point has remained poorly represented. The early detection obtained here reflects a finer sampling resolution and widens the temporal window in which the gene is known to be active. The detection at 3 DPA was obtained in whole fruit, a matrix that includes non-expressing tissue, and therefore constitutes a minimum estimate of the actual abundance in the placenta.

The temporal pattern of VAMT expression coincides with the period of active capsaicinoid accumulation (Arce-Rodríguez & Ochoa-Alejo, 2019) and suggests that events after fertilization participate in the establishment of the pathway in placental tissue, although genotype and environmental conditions may also modify the timing of activation (Kusaka et al., 2024).

### 2. Spray delivery achieves injection-level *VAMT* silencing

The capacity of dsR*VAMT* was validated by direct injection into the fruit pedicel. With the highest dose, an 89.5 % reduction in transcript abundance was obtained, comparable to that reported for other methods that also generate a localized disruption of the tissue: *VAMT* transcript was rendered undetectable by particle bombardment in developing fruits (Abraham-Juárez et al., 2008), and dsRNA targeting *EPSPS* was applied to amaranth leaves after abrasion with a carborundum solution (Sammons et al., 2011). What these antecedents share is that the surface barrier was bypassed by physical means.

Spraying was evaluated within the same comparative trial. Injected dsR*VAMT* reduced *VAMT* transcript abundance by 90.6 % and sprayed dsR*VAMT* by 86.2 %, with no significant differences between the two routes. Since both treatments were derived from the same plants, on the same date, and with the same normalization, the equivalence observed cannot be attributed to variation between trials. It is therefore indicated that, under the conditions evaluated, the efficacy of dsR*VAMT* was not delimited by the tissue penetration barrier, which carries agronomic implications, since spraying is scalable to extensive crops, unlike injection or particle bombardment, which operate on a plant-by-plant basis. The viability of this route at commercial scale is already supported by a regulatory precedent: Calantha, a foliar-applied dsRNA bioinsecticide directed against *Leptinotarsa decemlineata* in potato, was the first sprayable dsRNA product to be registered with the United States Environmental Protection Agency (Narva et al., 2025).

It is proposed that the success of the application was related to the availability of the stomatal route at the time of treatment. Although no foliar dsRNA uptake pathway has been conclusively confirmed, stomata have been suggested as an entry point, and it has been considered reasonable that sprayed dsRNA enters primarily through larger openings such as the stomata, the ectodesmata, or the hydathodes (Hoang et al., 2022). In the opposite direction, it has been documented that the physiological conditions at the time of application modify the efficacy of exogenously induced silencing, and that application at later periods of the day, particularly at night, is more efficient than daytime application, which suggests a link between plant circadian rhythms and the cellular machinery responsible for exoRNAi induction (Kiselev et al., 2021b).

The two observations have not been reconciled: the first predicts greater internalization during the day, when the stomata are open, whereas the second places the optimum in the night period, so that the determinants of dsRNA internalization remain an open field. In the present work, spraying was performed at 08:00 h, whereas in the work of Suprun et al. (2023), application was carried out between 21:00 and 21:30 h. The magnitude of inhibition obtained in the morning schedule is consistent with the stomatal route as the predominant entry pathway under the conditions evaluated, although the time of application was not examined as an experimental variable in this work.

In future work, delivery vehicles that prolong dsRNA availability on the treated surface should be compared. It has been demonstrated that loading dsRNA onto layered double hydroxide nanosheets protects it from nuclease degradation and from washing off and allows it to be detected on the leaf 30 days after a single spray, as opposed to the 20 days at which naked dsRNA becomes almost undetectable (Mitter et al., 2017). Increased application frequency from 3–5 DPA onwards and quantification of silencing in non-treated tissues should likewise be evaluated.

### 3. Silencing is modulable by dose

The three injected doses produced significantly different silencing levels for both *VAMT* and *Pun1*, with the six pairwise comparisons significant in both genes. The reduction in *VAMT* transcript was 26.0 % with 10 pmol, 36.2 % with 100 pmol, and 89.5 % with 1000 pmol, equivalent to 0.135, 1.35, and 13.5 µg of duplex per fruit. The response therefore does not constitute a trend, but a relationship distinguishable across the three levels evaluated. For reference, 70 µg per plant were employed in a single application for the silencing of four *MYB* repressors in tomato leaves, where the reduction in mRNA was not quantified (Suprun et al., 2023). In potato, modification of the tuber phenotype required six sprays of dsRNA targeting the isoamylases ISA1, ISA2, and ISA3 over fifteen weeks (Simon et al., 2023).

### 4. The response of *Pun1* to *VAMT* silencing

*Pun1* transcript abundance decreased in a dose-dependent manner in the injection experiment. This co-regulation is consistent with the association reported between both genes across cultivars, where high capsaicinoid accumulation is accompanied by high expression of both *VAMT* and *Pun1*, whereas both remain low in non-pungent varieties (Ogawa et al., 2015). The co-regulation between nodes of the pathway has also been documented in the reverse direction: silencing of the *AT3* gene, corresponding to the *Pun1* locus, by virus-induced gene silencing, reduced the expression of *VAMT* in chili pepper fruits (Arce-Rodríguez & Ochoa-Alejo, 2015). The reciprocity is thereby completed, since the silencing of *VAMT* was accompanied by a reduction of *Pun1*. The present results extend that association from a comparison between genotypes to an induced response within a single genotype, since the reduction of *Pun1* was obtained transiently and in a dose-dependent manner after silencing of the preceding node. A direct off-target effect does not explain the observation: alignment of the *Pun1* coding sequence (LC423556) against siRVAMT and its complementary strand, under default parameters, yielded no pairings, which rules out direct interaction, although it does not exclude propagation of silencing through secondary pathways.

Wounding has been reported to modify the expression of capsaicinoid biosynthetic genes, including *Pun1* and *pAMT* (Yang et al., 2024), which could contribute to the response observed in the injection experiment. However, *Pun1* abundance also decreased under spray application, where no wounding occurs, which indicates that the reduction is not solely a wound response. Capsinoid esters were nonetheless detected in the treated group even though the abundance of the *Pun1* transcript was reduced by approximately half.

### 5. Redirection of the vanillylamine branch point

The seven vanilloid compounds detected showed a consistent pattern with the blockade of the node. The five vanillylamine-derived amides decreased or ceased to be detectable, whereas the two capsinoid esters, absent in control, were detected in the treated group. The concordance among compounds is a more robust support than the decline of a single metabolite, since all of them share the same intermediate. Simple spraying of naked dsRNA or siRNA in water, without high pressure, abrasion, or a carrier, has repeatedly failed to produce detectable silencing (Dalakouras et al., 2016; Uslu et al., 2020), so the changes observed are attributable to dsRVAMT rather than to the aqueous vehicle.

This pattern is consistent with the mechanism described for *VAMT* loss of function: in the absence of vanillylamine, *Pun1* condenses vanillyl alcohol with fatty acids and capsinoids accumulate instead of capsaicinoids (Lang et al., 2009; Tanaka, 2025; Fig. 6). The magnitude of the change allows a direct comparison. In the CH-19 Sweet mutant, total capsaicinoids are reduced by 90.8 % and capsinoids, nearly absent in the pungent line, become the dominant fraction (Lang et al., 2009). In the present work, the summed concentration of the five vanillylamine-derived amides was reduced by 81.9 % and the capsinoid esters, non-detectable in the control, were detected in the treated group. The direction of the change is the same, and its magnitude is smaller, which is congruent with a transient silencing of 86.2 % as opposed to a complete and inherited loss of function. It is therefore indicated that the treatment partially reproduced the VAMT mutant phenotype via the topical route and without genetic modification.

**Fig. 6.**
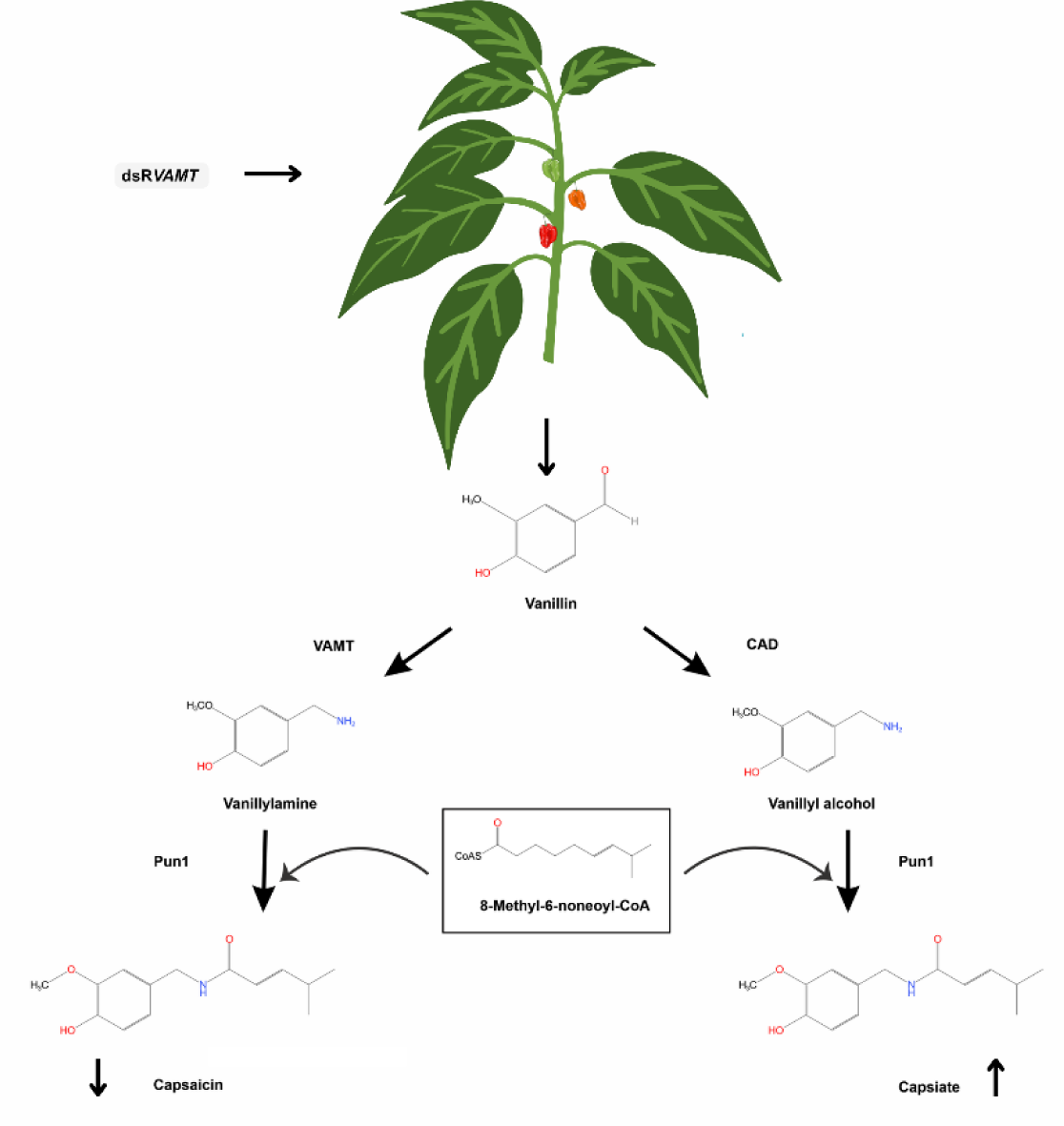
Schematic representation of dsRNA-mediated *VAMT* silencing and metabolic redirection of the vanillylamine branch point. Exogenous application of dsRNA leads to siRNA processing and subsequent inhibition of *VAMT*, decreasing capsaicinoids accumulation and redirecting intermediate flux toward vanillyl alcohol conversion, and induces capsinoid synthesis through *Pun1* activity

Variability among individual fruits provides an internal interpretation: the replicate that retained the highest capsaicin and dihydrocapsaicin concentrations was the same one in which neither of the two esters was detected, whereas the two replicates with the lowest amide content were those in which the esters appeared (Table S5). It is suggested that the decline of amides and the appearance of esters therefore behaved as manifestations of a single response. This heterogeneity suggests that the effect of SIGS-mediated silencing was not uniform among individual fruits. Its characterization requires a greater number of biological replicates and, given that variability could arise from differences in dsRNA retention on the fruit, a comparison of delivery vehicles with distinct adherence and release capacities.

The detection of both esters in the treated group suggests that repeated application throughout fruit development could accumulate capsinoids at levels of agronomic interest, given that these analogues lack pungency and retain functional activity. Nevertheless, a complete evaluation would require a scheme of successive applications and the harvest of fruits at commercial maturity, in addition to the quantification of capsinoids by a chromatographic method with an internal standard, to assess possible losses or degradation of the analyte during the extraction procedure.

### 6. Transport of siR*VAMT* to non-treated tissues

siRVAMT was detected in non-treated flowers and leaves of injected plants, and was not detected in the same tissues of untreated plants. Since the application was directed at the fruit while ensuring that the rest of the plant was not wetted, the presence of the siRNA in distant organs corresponds to transport and not to direct contact. This result is congruent with the systemic movement of exogenously applied dsRNA described in peanut (Faustinelli et al., 2018), canola (Willow et al., 2023), and barley (Biedenkopf et al., 2020).

Silencing in those distal tissues was nevertheless not quantified. That measurement was not part of the experimental design, which was oriented towards evaluating the local effect of spraying. Systemic detection therefore opens the possibility of regulating capsaicinoid accumulation in fruits other than the treated one, which would broaden the agronomic scope of the method. Its evaluation requires a design in which fruits from the same plant are protected during each application and subsequently analysed, so that silencing in non-exposed tissue can be quantified unambiguously. Likewise, these findings position the reproductive tissues as a compelling starting point for future work, particularly regarding whether siRVAMT exerts transgenerational effects through seed and offspring, a question that now warrants dedicated attention.

## Conclusions

*VAMT* transcript abundance was reduced, after a single spray of dsRVAMT applied without prior tissue abrasion, by a magnitude statistically indistinguishable from that obtained by injection, and the degree of inhibition was found to be adjustable through the applied dose. Inhibition of the metabolic node was accompanied by a reduction in capsaicin accumulation and by the appearance of capsinoid esters absent from control fruits, a pattern consistent with a redirection of flux at the vanillylamine branch point. Through the co-reduction of *Pun1*, an association previously described only across cultivars was reproduced within a single genotype and transiently. siRVAMT was detected in non-treated organs, and the reach of the effect beyond the treated tissue and the extension of silencing to other nodes of the pathway are left as open questions.

## Statements and Declarations

### Author Contributions

Conceptualization: C.C., R.G.G.G.; Methodology: E.A.-O., A.K.R.-C., R.G.G.G., C.C.; Formal analysis: E.A.-

O., C.C.; Investigation: E.A.-O., A.K.R.-C.; Visualization: E.A.-O.; Writing – original draft: E.A.-O., C.C.;

Writing – review & editing: A.K.R.-C., I.T.-P., R.V.O.V., R.G.G.G., C.C.; Supervision: C.C.; Funding acquisition: R.V.O.V., C.C. All authors read and approved the final manuscript.

### Data Availability

The datasets generated and analysed during the current study are available from the corresponding author on reasonable request.

### Ethics approval

This study did not involve human participants, animals, or endangered plant species. The *Capsicum chinense* cultivar used is a commercially available crop.

### Electronic Supplementary Material

ESM_1 (.docx) Supplementary tables: nutrient formulation supplied by fertigation (Table S1), manufacturer’s technical data sheet of the surfactant (Table S2), primer sequences (Table S3), calibration and quantification parameters for GC-MS analysis (Table S4), and individual capsaicinoid and capsinoid concentrations in placental tissue (Table S5)

## References

Abraham-Juárez MR, Rocha-Granados MDC, López MG, Rivera-Bustamante RF, Ochoa-Alejo N (2008) Virus-induced silencing of Comt, pAmt and Kas genes results in a reduction of capsaicinoid accumulation in chilli pepper fruits. Planta 227:681–695. 10.1007/s00425-007-0651-7

Arce-Rodríguez ML, Ochoa-Alejo N (2015) Silencing AT3 gene reduces the expression of pAmt, BCAT, Kas, and Acl genes involved in capsaicinoid biosynthesis in chili pepper fruits. Biol Plant 59:477–484. 10.1007/s10535-015-0525-y

Arce-Rodríguez ML, Ochoa-Alejo N (2019) Biochemistry and molecular biology of capsaicinoid biosynthesis: recent advances and perspectives. Plant Cell Rep 38:1017–1030. 10.1007/s00299-019-02406-0

Aza-González C, Núñez-Palenius HG, Ochoa-Alejo N (2011) Molecular biology of capsaicinoid biosynthesis in chili pepper (Capsicum spp.). Plant Cell Rep 30:695–706. 10.1007/s00299-010-0968-8

Biedenkopf D, Will T, Knauer T, Jelonek L, Furch ACU, Busche T, Koch A (2020) Systemic spreading of exogenous applied RNA biopesticides in the crop plant Hordeum vulgare. ExRNA 2:12. 10.1186/s41544-020-00052-3

Cedillo-Jiménez CA, Guevara-González RG, Cruz-Hernández A (2024) Exogenous dsRNA sequence based on miR1917 downregulates its target gene related to ethylene signaling in tomato seedlings and fruit. Sci Hortic 331:113090. 10.1016/j.scienta.2024.113090

Chen N, Dai X, Hu Q, Tan H, Qiao L, Lu L (2023) Sprayable double-stranded RNA mediated RNA interference reduced enzymatic browning of fresh-cut potatoes. Postharvest Biol Technol 206:112563. 10.1016/j.postharvbio.2023.112563

Cisneros AE, Carbonell A (2022) Systemic silencing of an endogenous plant gene by two classes of mobile 21-nucleotide artificial small RNAs. Plant J 110:1166–1181. 10.1111/tpj.15730

Dai X, Zhao PX (2011) psRNATarget: a plant small RNA target analysis server. Nucleic Acids Res 39:W155–W159. 10.1093/nar/gkr319

Dai X, Zhuang Z, Zhao PX (2018) psRNATarget: a plant small RNA target analysis server (2017 release). Nucleic Acids Res 46:W49–W54. 10.1093/nar/gky316

Dalakouras A, Wassenegger M, McMillan JN, Cardoza V, Maegele I, Dadami E, Runne M, Krczal G, Wassenegger M (2016) Induction of silencing in plants by high-pressure spraying of in vitro-synthesized small RNAs. Front Plant Sci 7:1327. 10.3389/fpls.2016.01327

Dubrovina AS, Aleynova OA, Suprun AR, Ogneva ZV, Kiselev KV (2020) Transgene suppression in plants by foliar application of in vitro-synthesized small interfering RNAs. Appl Microbiol Biotechnol 104:2125–2135. 10.1007/s00253-020-10355-y

Faustinelli PC, Power IL, Arias RS (2018) Detection of exogenous double-stranded RNA movement in in vitro peanut plants. Plant Biol 20:444–449. 10.1111/plb.12703

Ghosh P, Dullea R, Fischer JE, Turi TG, Sarver RW, Zhang C, Basu K, Das SK, Poland BW (2009) Comparing 2-nt 3′ overhangs against blunt-ended siRNAs: a systems biology based study. BMC Genomics 10(Suppl 1):S17. 10.1186/1471-2164-10-S1-S17

Hoang BT, Fletcher SJ, Brosnan CA, Ghodke AB, Manzie N, Mitter N (2022) RNAi as a foliar spray: efficiency and challenges to field applications. Int J Mol Sci 23:6639. 10.3390/ijms23126639

Kiselev KV, Suprun AR, Aleynova OA, Ogneva ZV, Kalachev AV, Dubrovina AS (2021a) External dsRNA downregulates anthocyanin biosynthesis-related genes and affects anthocyanin accumulation in Arabidopsis thaliana. Int J Mol Sci 22:6749. 10.3390/ijms22136749

Kiselev KV, Suprun AR, Aleynova OA, Ogneva ZV, Dubrovina AS (2021b) Physiological conditions and dsRNA application approaches for exogenously induced RNA interference in Arabidopsis thaliana. Plants 10:264. 10.3390/plants10020264

Koch A, Biedenkopf D, Furch A, Weber L, Rossbach O, Abdellatef E et al (2016) An RNAi-based control of Fusarium graminearum infections through spraying of long dsRNAs involves a plant passage and is controlled by the fungal silencing machinery. PLoS Pathog 12:e1005901. 10.1371/journal.ppat.1005901

Kusaka H, Nakasato S, Sano K, Kobata K, Ohno S, Doi M, Tanaka Y (2024) An evolutionary view of vanillylamine synthase pAMT, a key enzyme of capsaicinoid biosynthesis pathway in chili pepper. Plant J 117:1453–1465. 10.1111/tpj.16573

Lang Y, Kisaka H, Sugiyama R, Nomura K, Morita A, Watanabe T, Tanaka Y, Yazawa S, Miwa T (2009) Functional loss of pAMT results in biosynthesis of capsinoids, capsaicinoid analogs, in Capsicum annuum cv. CH-19 Sweet. Plant J 59:953–961. 10.1111/j.1365-313X.2009.03921.x

Lück S, Kreszies T, Strickert M, Schweizer P, Kuhlmann M, Douchkov D (2019) siRNA-Finder (si-Fi) software for RNAi-target design and off-target prediction. Front Plant Sci 10:1023. 10.3389/fpls.2019.01023

Malm L, Palm E, Souihi A, Plassmann M, Liigand J, Kruve A (2021) Guide to semi-quantitative non-targeted screening using LC/ESI/HRMS. Molecules 26:3524. 10.3390/molecules26123524

Mitter N, Worrall EA, Robinson KE, Li P, Jain RG, Taochy C, Fletcher SJ, Carroll BJ, Lu GQ, Xu ZP (2017) Clay nanosheets for topical delivery of RNAi for sustained protection against plant viruses. Nat Plants 3:16207. 10.1038/nplants.2016.207

Mückstein U, Tafer H, Hackermüller J, Bernhart SH, Stadler PF, Hofacker IL (2006) Thermodynamics of RNA–RNA binding. Bioinformatics 22:1177–1182. 10.1093/bioinformatics/btl024

Nakaniwa R, Misawa Y, Nakasato S, Sano K, Tanaka Y, Nakatani S, Kobata K (2024) Biochemical aspects of putative aminotransferase responsible for converting vanillin to vanillylamine in the capsaicinoid biosynthesis pathway in Capsicum plants. J Agric Food Chem 72:559–565. 10.1021/acs.jafc.3c07369

Narva K, Otto E, Sridharan K, Flannagan R, Barnes E, Mézin L, Manley B (2025) Calantha™: the first commercialized sprayable dsRNA product for insect control. In: RNA interference in agriculture: basic science to applications. Springer, Cham, pp 679–715. 10.1007/978-3-031-81549-2_27

Nityagovsky NN, Kiselev KV, Suprun AR, Dubrovina AS (2022) Exogenous dsRNA induces RNA interference of a chalcone synthase gene in Arabidopsis thaliana. Int J Mol Sci 23:5325. 10.3390/ijms23105325

Ogawa K, Murota K, Shimura H et al (2015) Evidence of capsaicin synthase activity of the Pun1-encoded protein and its role as a determinant of capsaicinoid accumulation in pepper. BMC Plant Biol 15:93. 10.1186/s12870-015-0476-7

Reddy DN, Mylabathula MM, Al-Rajab AJ (2024) Industrial demand and applications of capsaicin. In: Swamy MK, Kumar A (eds) Capsaicinoids. Springer, Singapore. 10.1007/978-981-99-7779-6_12

Sammons R, Ivashuta S, Liu H, Wang D, Feng P, Kouranov A, Andersen S (2011) Polynucleotide molecules for gene regulation in plants. US Patent 20110296556 A1

Sano K, Uzawa Y, Kaneshima I, Nakasato S, Hashimoto M, Tanaka Y, Nakatani S, Kobata K (2022) Vanillin reduction in the biosynthetic pathway of capsiate, a non-pungent component of Capsicum fruits, is catalyzed by cinnamyl alcohol dehydrogenase. Sci Rep 12:12384. 10.1038/s41598-022-16150-1

Simon I, Persky Z, Avital A, Harat H, Schroeder A, Shoseyov O (2023) Foliar application of dsRNA targeting endogenous potato (Solanum tuberosum) isoamylase genes ISA1, ISA2, and ISA3 confers transgenic phenotype. Int J Mol Sci 24:190. 10.3390/ijms24010190

Suprun AR, Kiselev KV, Dubrovina AS (2023) Exogenously induced silencing of four MYB transcription repressor genes and activation of anthocyanin accumulation in Solanum lycopersicum. Int J Mol Sci 24:9344. 10.3390/ijms24119344

Szabat M, Kierzek R (2017) Parallel-stranded DNA and RNA duplexes – structural features and potential applications. FEBS J 284:3986–3998. 10.1111/febs.14187

Tanaka Y (2025) Recent understanding of the biosynthesis of capsaicinoids and low-pungent analogs towards quality improvement of chili pepper. Hortic J 94:117–128. 10.2503/hortj.SZD-R002

Uslu VV, Bassler A, Krczal G, Wassenegger M (2020) High-pressure-sprayed double stranded RNA does not induce RNA interference of a reporter gene. Front Plant Sci 11:534391. 10.3389/fpls.2020.534391

Varkonyi-Gasic E, Wu R, Wood M, Walton EF, Hellens RP (2007) Protocol: a highly sensitive RT-PCR method for detection and quantification of microRNAs. Plant Methods 3:12. 10.1186/1746-4811-3-12

Warnock ND, Wilson L, Canet-Perez JV, Fleming T, Fleming CC, Maule AG, Dalzell JJ (2016) Exogenous RNA interference exposes contrasting roles for sugar exudation in host-finding by plant pathogens. Int J Parasitol 46:473–477. 10.1016/j.ijpara.2016.02.005

Willow J, Kallavus T, Soonvald L, Caby F, Silva AI, Sulg S, Kaasik R, Veromann E (2023) Examining spray-induced gene silencing for pollen beetle control. J Nat Pestic Res 5:100036. 10.1016/j.napere.2023.100036

Yang Y, Gao C, Ye Q, Liu C, Wan H, Ruan M, Zhou G, Wang R, Li Z, Diao M, Cheng Y (2024) The Influence of Different Factors on the Metabolism of Capsaicinoids in Pepper (Capsicum annuum L.). Plants 13:2887. 10.3390/plants13202887

Zhang ZX, Zhao SN, Liu GF, Huang ZM, Cao ZM, Cheng SH, Lin SS (2016) Discovery of putative capsaicin biosynthetic genes by RNA-Seq and digital gene expression analysis of pepper. Sci Rep 6:34121. 10.1038/srep34121

