## Supplemental Table 1-5 for "A single dsRNA spray silences *VAMT* and shifts habanero pepper fruit metabolism towards capsinoids"

Electronic Supplementary Material

Plant Cell Reports

Estefania Arellano-Ordoñez, Amanda Kim Rico-Chávez, Irineo Torres-Pacheco, Rosalia Virginia Ocampo-Velázquez, Ramón Gerardo Guevara-González, Christopher Cedillo

**Table S1. Nutrient formulation supplied by fertigation at each phenological stage.**

| Nutrient | Vegetative stage | Flowering stage | Fruit set stage | Production stage |
|---|---|---|---|---|
| $NO_3^-$ | 536 | 942 | 496 | 682 |
| $H_2PO_4^-$ | 8.4 | 82 | 165 | 343 |
| $SO_4^{2-}$ | 554 | 402 | 235 | 273 |
| $Ca^{2+}$ | 199 | 228 | 120 | 160 |
| $Mg^{2+}$ | 52 | 42 | 23 | 28 |
| $K^+$ | 258 | 258 | 175 | 262 |
| $Fe^{2+}$ | 2 | 2 | 2 | 4 |
| $Zn^{2+}$ | 0.05 | 0.05 | 0.05 | 0.05 |
| $Mn^{2+}$ | 0.21 | 0.21 | 0.21 | 1 |
| $Cu^{2+}$ | 0.5 | 0.5 | 0.5 | 0.5 |

Note: Concentrations are expressed as mg $L^{-1}$. Micronutrients were supplied in chelated form.

**Supplementary Table S2. AGRONATURALIA manufacturer's technical data sheet**

| Category | Parameter / Component | Description / Value |
|---|---|---|
| Product Identification | Brand name | EcoNano MegaKliner |
| | Purpose and substance type | Commercial, industrial, and agricultural surfactant, degreaser, and adjuvant |
| | Regulatory status | U.S. EPA FIFRA registration not required |
| Physicochemical Properties | Physical form and appearance | Flowable viscous liquid; clear to honey-colored |
| | Odor | Nondescript |
| | Specific gravity | 1.001 (≈ 8.338 lbs./gal.) |
| | Potential of hydrogen (pH) | 9.2 |
| | Vapor pressure | Not established (equivalent to water) |
| | Flash point | None (aqueous-based, non-flammable product) |
| | Operational temperature limits | Do not expose to temperatures below 40°F (4.4°C) or above 130°F (54.4°C) |
| Composition & Potential Ingredients | Organic solvents and acids | Organic alcohol (CAS 97-99-4), Ethyl lactate (CAS 97-64-3), Gluconic acid (CAS 526-95-4), Water (CAS 007732-18-5) |
| | Surfactants | Alkyl polyglycoside (CAS 141464-42-8), Coco polyglycoside (CAS 151911-53-4), Alkanolamines (CAS 141-43-5), Tall oil fatty acids (CAS 61790-12-3) |
| | Chelating agents | EDDS (CAS 20846-91-7), Disodium EDTA (Kosher) (CAS 6381-92-6) |
| | Botanical oils and extracts | Almond oil (CAS 8007-69-0), Cedar oil (CAS 8000-27-9), Cinnamon oil (CAS 8007-80-5), Corn oil (CAS 8001-30-7), Geranium oil (CAS 8000-46-2), Jojoba oil (CAS 61789-91-1), Lemongrass oil (CAS 8007-02-1), Soybean oil (CAS 8001-22-7), Processed sugar cane extracts (CAS 57-50-1) |

| | | |
|---|---|---|
| Toxicity and Safety Profile | Carcinogenicity | Contains no known carcinogens |
| | Acute toxicity (Lethal Dose) | $LD_{50}$ Oral: $> 1500$ mg/kg; $LD_{50}$ Dermal: $> 2000$ mg/kg |
| | Ocular effects | MMTS (Maximum Mean Total Score) between 15 and 25; causes mild, reversible soapy irritation |
| Stability and Environmental Fate | Chemical stability | Stable; hazardous polymerization will not occur |
| | Biodegradability | Aqueous-based, readily biodegradable product |
| | Decomposition products | None known |
| Transportation and Handling | D.O.T. Shipping Name | Commercial, industrial, and agricultural non-hazardous surfactant |
| | D.O.T. Hazard Class / Labels | Not established (N/E); no labels required |
| | Shelf life | One year (when stored at normal ambient room or warehouse conditions) |

**Table S3. Sequences of the primers used.**

| **Gene** | **Accession number** | **Orientatio n** | **Sequence (5′→3′)** | **Source** |
|---|---|---|---|---|
| *VAMT* | LC423555 | Forward | CCACTTACATTTCTGCTGGTCTCTC | Sano et al. (2022) |
| | | Reverse | CAATGAAAGCAGCTACTGTTTCAGG | Sano et al. (2022) |
| *Pun1* | LC423556 | Forward | GCCTTGGGCGAATAATTGTGAAG | Sano et al. (2022) |

| | | | | |
|---|---|---|---|---|
| | | Reverse | TTAAGCAGAGAGCAACCATCACC | Sano et al. (2022) |
| *β-actin* | AY572427 | Forward | AGCAACTGGGACGATATGGAGAAG | Sano et al. (2022) |
| | | Reverse | AAGAGACAACACCGCCTGAATAGC | Sano et al. (2022) |
| siR*VAMT* | - | Stem-loop RT primer | GTTGGCTCGGTGCAGGGTCCGAGGTATTCGCACCAGAGCCACACTTTC | This work |
| | | Specific forward | TTCACAAACTCTGTAGAAAG | This work |
| | | Universal reverse | GTGCAGGGTCCGAGGT | This work |

**Table S4. Calibration and quantification parameters for GC-MS analysis.**

| Analyte | RT (min) | Range (µg/mL) | n | $R^2$ | LOD (µg/mL) | LOQ (µg/mL) | Calibration curve applied |
|---|---|---|---|---|---|---|---|
| Capsaicinoids | | | | | | | |
| Nonivamide | 20.4 | - | - | - | 0.0116 | 0.0388 | Capsaicin (surrogate) |
| Putative amide | 21.1 | - | - | - | 0.0116 | 0.0388 | Capsaicin (surrogate) |
| Capsaicin | 21.8 | 0–300 | 7 | 0.9938 | 0.0116 | 0.0388 | Own standard |
| Dihydrocapsaicin | 22.2 | 0–300 | 7 | 0.9915 | 0.0464 | 0.1548 | Own standard |
| Homodihydrocapsaicin | 24.1 | - | - | - | 0.0464 | 0.1548 | Dihydrocapsaicin (surrogate) |
| Capsinoids | | | | | | | |
| Capsiate | 17.3 | 0–1500 | 6 | 0.8994 | 0.3039 | 1.0129 | Own standard |

| | | | | | | | |
|---|---|---|---|---|---|---|---|
| Capsinoid | 17.5 | - | - | - | 0.3039 | 1.0129 | Capsiate (surrogate) |

Note a. Calibration standards were injected in split mode (5:1) and samples in splitless mode; the split ratio was applied as a dilution factor to the concentrations obtained for the samples. All values in this table are expressed on the split-corrected scale.

Note b. Calibration standards were injected in July 2024 and samples in December 2025. Successive trimming of the column during that interval displaced retention times by approximately 0.2 min; the retention times reported correspond to the sample runs.

Note c. LOD and LOQ were determined from the signal-to-noise ratio, at ratios of 3:1 and 10:1, respectively. Compounds for which no authentic standard was available were semi-quantified against the curve indicated and inherited its limits. "Not detected" indicates a signal below the LOD and does not establish absence of the compound.

**Table S5.** Individual capsaicinoid and capsinoid concentrations in placental tissue of control and dsRVAMT SIGS-treated fruits.

| **Compound** | **RT (min)** | **Control 1** | **Control 2** | **Control 3** | **dsRNA 1** | **dsRNA 2** | **dsRNA 3** |
|---|---|---|---|---|---|---|---|
| Capsaicinoids | | | | | | | |
| Nonivamide | 20.4 | 3.01 | 1.88 | 2.74 | n.d. | n.d. | n.d. |
| Putative amide | 21.1 | 0.57 | 0.38 | 0.39 | n.d. | n.d. | n.d. |
| Capsaicin | 21.8 | 32.18 | 20.21 | 28.94 | 3.88 | 2.90 | 5.87 |
| Dihydrocapsaicin | 22.2 | 9.96 | 6.39 | 9.04 | 1.39 | 1.36 | 5.24 |
| Homodihydrocapsaicin | 24.1 | 0.66 | 0.38 | 0.42 | 0.03 | 0.05 | 0.53 |
| Total vanillylamine-derived amides | - | 46.38 | 29.24 | 41.53 | 5.30 | 4.31 | 11.64 |
| Capsinoids | | | | | | | |
| Capsiate | 17.3 | n.d. | n.d. | n.d. | 15.29 | 11.61 | n.d. |
| Capsinoid | 17.5 | n.d. | n.d. | n.d. | 2.66 | 4.08 | n.d. |

Note: Values are expressed as mg/g dry weight. n.d., not detected, that is, signal below the limit of detection. Retention times correspond to the sample runs. Capsaicin and dihydrocapsaicin were quantified against their own calibration curves; homodihydrocapsaicin was semi-quantified against the dihydrocapsaicin curve, and nonivamide and the putative amide against the capsaicin curve. The values for capsiate and the capsinoid were obtained from a curve that did not meet the adopted linearity criterion and are presented for transparency; in the manuscript both esters are reported as detection frequency. Compounds recorded as not detected were assigned a value of zero for the calculation of the sum.
